# Mobile microscopy and telemedicine platform assisted by deep learning for quantification of *Trichuris trichiura* infection

**DOI:** 10.1101/2021.01.19.426683

**Authors:** Elena Dacal, David Bermejo-Peláez, Lin Lin, Elisa Álamo, Daniel Cuadrado, Álvaro Martínez, Adriana Mousa, María Postigo, Alicia Soto, Endre Sukosd, Alexander Vladimirov, Charles Mwandawiro, Paul Gichuki, Nana Aba Williams, José Muñoz, Stella Kepha, Miguel Luengo-Oroz

## Abstract

Soil-transmitted helminths (STH) are the most prevalent pathogens among the group of neglected tropical diseases (NTDs). Kato-Katz technique is the diagnosis method recommended by WHO and although is generally more sensitive than other microscopic methods in high transmission settings, it often presents a decreased sensitivity in low transmission settings and it is labour intensive. Digitizing the samples could provide a solution which allows to store the samples in a digital database and perform remote analysis. Artificial intelligence methods based on digitized samples can support diagnostics efforts by support diagnostics efforts by performing an automatic and objective quantification of disease infection.

In this work, we propose an end-to-end pipeline for microscopy image digitization and automatic analysis of digitized images of soil-transmitted helminths. Our solution includes (1) a digitalization system based on a mobile app that digitizes the microscope samples using a low-cost 3D-printed microscope adapter, (2) a telemedicine platform for remote analysis and labelling and (3) novel deep learning algorithms for automatic assessment and quantification of parasitological infection of STH.

This work has been evaluated by comparing the STH quantification using both a manual remote analysis based on the digitized images and the AI-assisted quantification against the reference method based on conventional microscopy. The deep learning algorithm has been trained and tested on 41 slides of stool samples containing 949 eggs from 6 different subjects using a cross-validation strategy obtaining a mean precision of 98,44% and mean recall of 80,94%. The results also proved the potential of generalization capability of the method at identifying different types of helminth eggs.

In conclusion, this work has presented a comprehensive pipeline using smartphone-based microscopy integrated with a telemedicine platform for automatic image analysis and quantification of STH infection using artificial intelligence models.

## 1. Introduction

Soil-transmitted helminths (STH), which include hookworms (*Ancylostoma duodenale* and *Necator americanus*), roundworm (*Ascaris lumbricoides*) and whipworm (*Trichuris trichiura*) are the most prevalent pathogens among the group of neglected tropical diseases (NTDs) and predominantly occur in tropical and subtropical low- and middle-income countries (1). Globally STH affects over 1.5 billion individuals, causing a loss of more than 3 million disability adjusted life-years (DALYs) (2). The World Health Organization (WHO) 2030 Roadmap for NTDs set out strategies for STH control that focused on the elimination of this disease as a public health problem (3).

In many endemic countries, STH control strategy is implemented through targeted mass drug administration (MDA), using the anthelmintic drugs benzimidazoles (BZ) albendazole or mebendazole (4). Currently there are no reliable and cost effective diagnostic methods that can accurately evaluate the impact of the on-going MDA programmes. The diagnostic method recommended by WHO is Kato-Katz, a laboratory method for preparing human stool samples in a microscope smear using a small spatula and slide template that allows a standardized amount of faeces to be examined under a microscope and quantify STH infection (5,6). Kato-Katz, while being generally more sensitive than other microscopic methods in high transmission settings, requires limited equipment and is easy to perform in low resource settings (7). However, it often presents a decreased sensitivity in low transmission settings. One of the main disadvantages of the Kato-Katz technique is the necessity to read samples within 30 minutes of preparation as eggs tend to disappear or hatch, especially those of hookworms, and thus considerably reducing the sensitivity of this technique and even highly trained microscopists can misidentify species or give inconsistent results and even highly trained microscopists can misidentify species or give inconsistent results (8,9). Digitizing the samples at the right moment could provide a solution, which allows digital storage of the sample images and further analysis at any moment in time. Different digitalization devices have been tested allowing remote diagnosis and second opinion (10). A further step that could save time, increase performance and remove subjectivity of microscopic techniques is the possibility of implementing artificial intelligence algorithms for the automatic detection and quantification of these parasites on digitized image samples. This would be a major advance in the diagnosis and control of these diseases and its implementation would not disrupt the laboratories normal workflow since the basis of diagnosis is still microscopy and the technique used is the Kato-Katz.

Artificial intelligence-based technologies are rapidly evolving into applicable medical solutions and are actually revolutionizing the field (11). However, only few studies have been put effort to fulfill rigorous regulation standards to be approved by regulatory institutions such as the FDA (12). Most of these approved technologies were developed for the fields of radiology, cardiology and internal medicine. However, AI systems have also the potential to be applied to enable a rapid and objective diagnosis of NTDs and to enable public health delivery in low- and middle-income countries. In this context an special effort has to be made for the application of AI methods in such diseases as has been recommended by the World Health Organization (WHO) (13).

Several approaches for computer-aided analysis of helminth eggs detection and classification using artificial intelligence have been investigated in the last years. Alva *et al*. proposed the use of hand-crafted features along with a multivariate logistic regression for intestinal parasites classification (14). Other notable recent deep learning-based approach used a large fecal database with over 1122 patients including 22440 images for the identification of visible components in feces, including blood and epithelial cells, as well as STH eggs, proposing the so-called FecalNet (15). This work proved the potential of the use of these methods for the automatic analysis of stool samples using conventional microscopy images. Holmstrom et al. proposed to acquire microscopy images with a portable scan connected to a laptop and used a two stage sequential algorithm where candidates previously proposed by the first algorithm are classified as any type of helminth egg, and obtained promising results despite their limited number of training samples (16). The use of deep learning-based object detection methods for the automatic analysis and detection of helminth eggs on images acquired with smartphone-compatible microscopy attachments has been already tested (17).This work achieved comparable sensitivity to standard microscopy when detecting *Ascaris* spp. but showed a low performance in the identification of *Trichuris* spp. This is probably caused by the use of a cheap smartphone-compatible microscopy attachment (a magnification endoscope; USB Video Class, UVC) where the light source comes from the same direction as the camera. This produces images with insufficient quality specially for *Trichuris* spp. which have thinner and more translucent membranes. Previous approaches might disrupt the usual laboratory workflow as they do not use conventional microscopes.

The objective of this study is to (a) propose and develop an end-to-end system for remote and automatic detection and quantification of soil-transmitted helminths (STH), primarily for the detection of *T. trichuris*, based on digitized microscopy images acquired with a 3D printed low-cost adapter and artificial intelligence methods, and (b) to assess the proposed approach by comparing it with the conventional manual strategy used as reference.

## 2. Material and methods

In this work, we present an entire end-to-end pipeline from stool sample collection to automatic analysis of the samples for the identification of STH parasites. **Figure 1** schematizes the proposed end-to-end workflow, where the samples collected and prepared from different subjects are digitized and uploaded to a telemedicine platform for remote analysis. Additionally, the digitized samples are used for training AI algorithms in order to be able to perform an automatic analysis for future incoming digitized samples. All the prepared samples are also analyzed during the process by conventional microscopy methods for comparative purposes.

**Figure 1.**
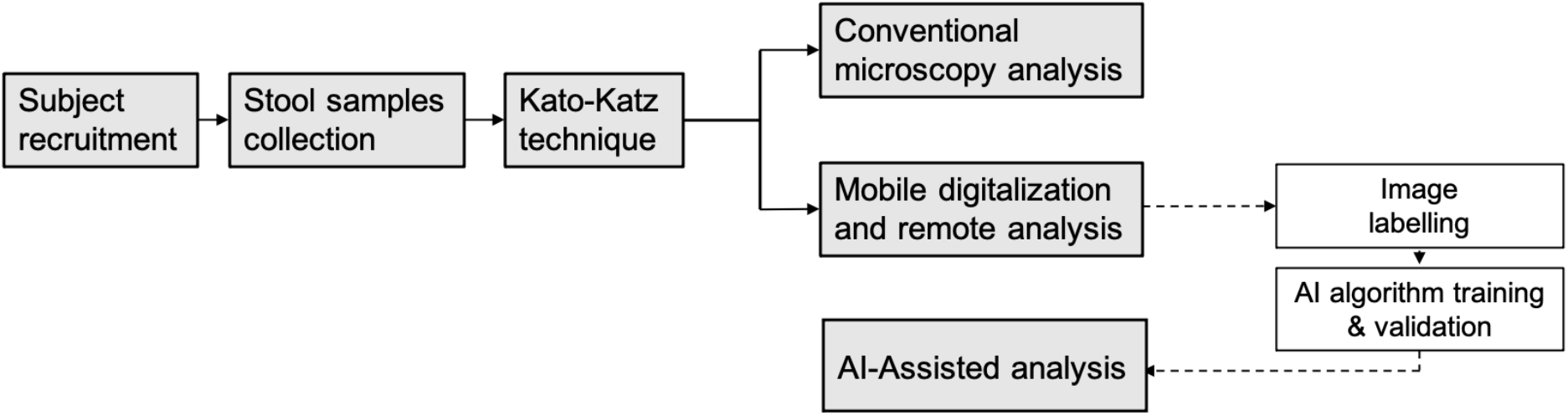
Schematic representation of study design and experimental workflow.

### 2.1. Data and Samples

Stool was collected from study participants from Kwale county who were part of a follow-up study related to exploring the persistence of *Trichuris trichiura* infection. All stool samples were transferred to the laboratory within four hours from collection, and preprocessed using Kato-Katz thick smear method (41.7mg template) and analyzed by conventional microscopy on the same day for identifying and quantifying the presence of STH eggs by 3 independent technicians. We digitized using the proposed digitalization pipeline samples from 12 subjects (6 positive and 6 negatives). These digitized samples were used for the evaluation of the remote analysis system as well as to train and evaluate the AI algorithm. From the 6 positive subjects, 5 were only positive for *Trichuris* spp. and one presented a co-infection of *Trichuris* spp. and *Ascaris* spp. For each positive stool sample, 7 slides were prepared, while for negative stool samples 1 or 2 slides were prepared. Ethical approval was obtained from the Kenya Medical Research Institute (KEMRI) Ethics Review Committee (SERU 3873).

### 2.2. Digitalization pipeline of clinical samples

The proposed digitalization system uses a low-cost 3D-printed device made with biodegradable material which allows coupling a mobile phone with a conventional optical microscope by aligning the smartphone camera with the microscope objective for acquiring images. The smartphone uses an Android mobile app specifically developed and customized for a fast and easy digitalization and sharing of digitized microscopy images (**Figure 2a**). Fifty-one Kato-Katz slides were digitized at 10x magnification using two different smartphone models (Xiaomi Pocophone F1 and Bq Aquaris X2) by attaching the 3D-printed device to the ocular of a conventional light microscope (Leica DM-2000). All microscopy fields where helminth eggs were present or suspected were digitized. In addition, visually-confirmed negative images were also acquired for both positive and negative subjects. Images were acquired in the JPG format with a resolution of 12 Mpx through the mobile application and uploaded via the mobile network to a cloud telemedicine platform.

**Figure 2.**
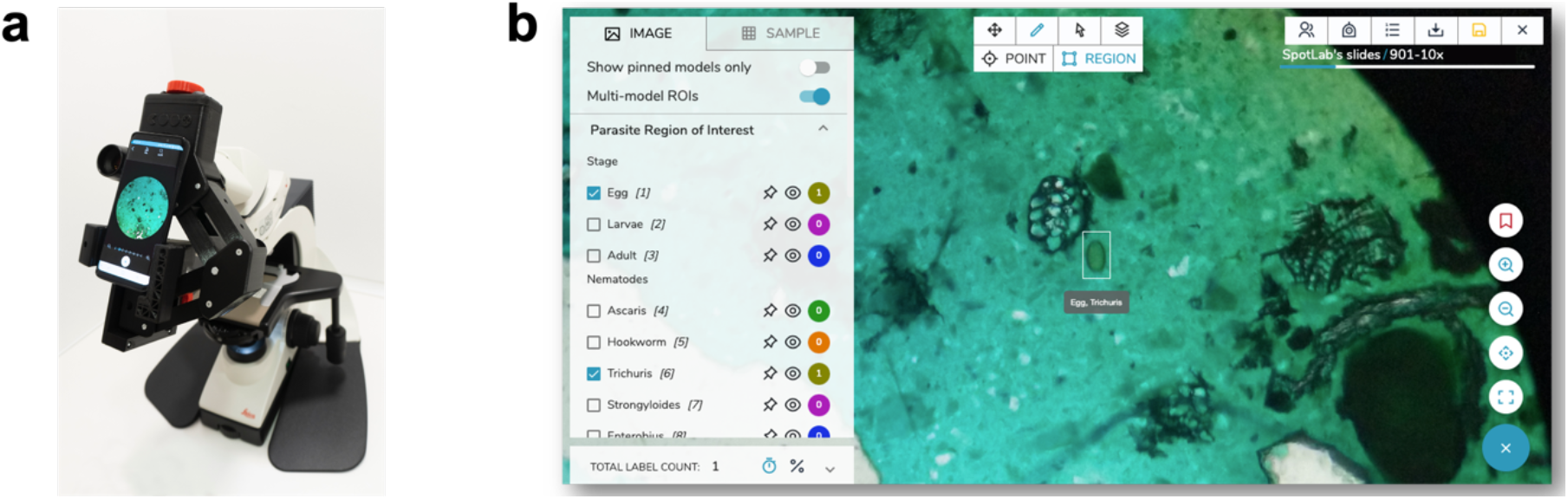
**a** 3D-printed adapter used for attaching a smartphone to a conventional microscope and digitizing samples through a mobile app. **b** Telemedicine web platform for viewing and remote analysis of the digitized samples performing a manual annotation.

### 2.3. Telemedicine platform for remote analysis

All acquired images are transferred from the smartphone to a telemedicine platform, where the images are stored and presented in an easy-to-use dashboard that allows its visualization, management and labelling (see **Figure 2b**). In this web platform, the standard clinical stic protocols are translated into digital tasks which are adapted to the clinical case and disease under study.

All acquired images were labelled through the telemedicine platform by one expert using a customized stool diagnostic protocol allowing to tag parasites that can be seen in the images and thus quantify parasitic infection. The image database together with the label data can be accessed by other professionals and coupled with other support platforms for diagnostic assistance.

### 2.4. Artificial intelligence algorithm

#### 2.4.1. Network architecture

The proposed approach for helminth eggs identification relies on a CNN-based detection algorithm. Most of CNN-based detection algorithms are designed to perform at the same time both object localization task, which determines where objects are located in a given image, and object classification task, which determines which particular category each previously located object belongs to.

In this paper, Single-Shot Detection (SSD) architecture (18) together with the backbone MobileNet network (19) for feature extraction were used. The model was initialized with pretrained weights on the COCO Image database (20). We used RMSprop optimizer with an exponential decay learning rate for minimizing the total loss function which was computed as the sum of a sigmoid cross entropy loss for object classification and smooth L1 loss for object localization. We also employed early stopping technique, where the training finishes before overfitting begins by stopping the training process when the error on the validation set does not decrease for a predefined number of steps.

#### 2.4.2. Training dataset generation

The training dataset was generated by extracting 512×512 pixels image patches around the location of labels which were placed manually by the experts on helminths. The size of the image patches was selected based on a balance between the relative size of the objects in relation to the size of image patches and the computational cost needed at inference time, as a small size of image patches increases the number of patches to be processed at test time using the sliding window procedure (see Section 3.3.4).

Additionally, and with the purpose of augmenting the training dataset size, we randomly selected 512×512 pixels image patches around the manually placed bounding boxes multiple times always ensuring that all labeled objects were fully covered. This was done so that each object can appear in different locations within the image patch and different context environments are captured. Moreover additional augmentation was conducted by applying on-the-fly random flip and rotation transformations during training.

The framework used for training the proposed method was based on Tensorflow Object Detection API using a cloud computing environment with a GPU Nvidia Tesla K80 12GB. The time needed for training the algorithm using the described hardware was approximately 3 hours.

## 3. Experiments and results

We established the conventional microscopy procedure performed by three different experts as the reference method. The reference measurement was computed as the average number of eggs counted by the three experts in each image field.

To evaluate and validate the entire proposed pipeline for digitizing and automatic assessing microscopy images, we analyzed the available database using the two described methodologies, i.e manually analysing digitized images through a remote telemedicine platform and using the proposed AI algorithm. The remote analysis and the AI pipelines have been compared against the manual pipeline used as reference.

### 3.1. Remote analysis of digitized samples

The digitalization of the 51 Kato-Katz slides was carried out by two technicians. Bq Aquaris X2 smartphone was used for the digitalization of 41 slides, while the remaining 10 slides were digitized by Xiaomi Pocophone F1 smartphone.

From the 51 digitized slides, a total of 1508 image fields were digitized and uploaded to the telemedicine platform and analyzed by a different person than those who digitized the slides, resulting in 797 positive images for at least one soil-transmitted helminth and 711 negative images. **Table 2** summarizes the digitized samples from all positive patients where a total of 949 *Trichuris* spp. egg labels were obtained. It should be noted that patient number 6 had a bi-parasite infection for *Trichuris* spp. and *Ascaris* spp., obtaining additional 4296 labels for *Ascaris* spp. eggs. Additionally, as expected, all 10 digitized slides from the 6 negative subjects obtained a negative result where no eggs were found in the images.

**Table 2.** Database of digitized Kato-Katz slide samples from 12 patients. *Patient number 5 had 6 slides instead of 7 because one of them broke during its handling.

|  | Patient | n° of slides | n° of (+) images | n° of (-) images | total number of images | n° *Trichuris* spp. Eggs |
| --- | --- | --- | --- | --- | --- | --- |
| **Positive subjects** | **1** | 7 | 94 | 39 | 133 | 148 |
|  | **2** | 7 | 39 | 15 | 54 | 47 |
|  | **3** | 7 | 50 | 21 | 71 | 74 |
|  | **4** | 7 | 28 | 203 | 231 | 30 |
|  | **5** | 6* | 44 | 10 | 54 | 47 |
|  | **6** | 7 | 542 | 13 | 555 | 603 |
| **Total** |  | **41** | **797** | **301** | **1098** | **949** |

The remote manual egg counting using digitized images was compared to the reference method based on the conventional microscopy procedure.

For the comparison of the two measurement methodologies (conventional microscopy and remote analysis of digitized samples) we excluded patient number 3 because its slides had not been well preserved and their reading could not be done correctly. To evaluate the correlation between the two egg measurements for each slide, Pearson correlation coefficient was calculated. The Pearson correlation coefficient was 0.95 (positive association) with a 95% confidence interval of 0.91−0.98 and a p-value <0.001 (p-value < 2.2e−16). In addition, the regression linear model was estimated to bey = 1,07× − 0,94 (see **Figure 3a**). These results indicate that both methods perform similarly, although the remote analysis of digitized samples slightly overestimates the results obtained with conventional microscopy.

**Figure 3.**
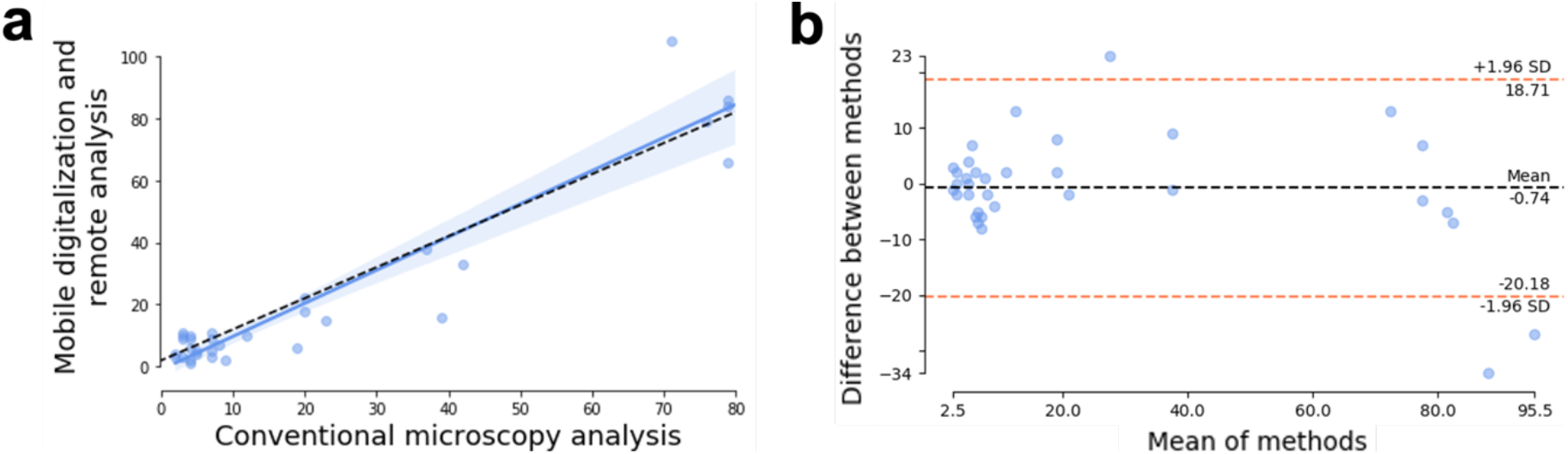
**a** Scatterplot of the manual counting using conventional microscopy vs. the remote counting of digitized samples. **b** Bland-Altman plot showing the difference between the conventional microscopy-based analysis and the remote analysis of digitized samples.

Furthermore, since a high correlation is not necessarily synonymous with agreement between methods, since it evaluates the relationship and not the difference, we calculated the agreement using the Bland-Altman method (21). **Figure 3b** shows the lower and upper limits (red dashed lines, with values of −18.71 and 20.18 respectively), and the mean bias (black dashed line) which took a value of −0.74 meaning that that the remote analysis of digitized samples measured 0.74 units more than the conventional microscopy method. These results show a good agreement between both methods.

Similar results were obtained when compared both methods at patient level. In this case, the Pearson correlation coefficient was 0.99 (positive association) with a 95% confidence interval of 0.82-0.99 and a p-value <0.01 (p-value = 0.002) and the Bland-Altman bias took a value of −0.86 indicating that the remote analysis of digitized samples measures in mean 0.86 units more than the conventional microscopy method, and thus reaffirming the good agreement between methods.

This slight overestimation of egg count observed in the remote analysis compared to the conventional methodology may indicate that when the count is performed remotely, the expert can perform a more exhaustive job being able to detect more eggs than when it is done in the field.

### 3.2. Artificial intelligence-assisted analysis

#### 3.2.1. Analysis of the model performance

To evaluate the proposed method for *Trichuris* spp. detection, we constructed a leave-one-out cross validation scheme at the patient level. Thus, the algorithm is applied and tested only on samples belonging to a single subject, using all other subject samples as a training set. This process ensures the test set is completely independent from the training set. It should be noted that as the proposed algorithm was focused on the detection of *Trichuris* spp., the leave-one-out scheme was constructed based only on those subjects that were positive for *Trichuris* spp.

To assess the model performance, we considered the precision (P), recall (R) and F-score (F) metrics defined as:

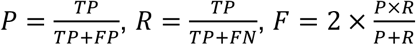

where TP, FP and FN denote the number of true positives, false positives and false negatives respectively. TP were defined as both correctly boxed and classified object, false detection was considered as FP and FN was defined as all ground truth objects misdetected by the algorithm or proposed for a wrong label. All proposed boxes by the algorithm had to have a certainty greater than or equal to 30% to be considered as proposed. The certainty of an object was defined based on the probability given by the algorithm and associated to the predicted label of this object. Correctly proposed and classified boxes were considered as true positive detections when had an intersection over union (IOU) with the ground truth of 30%. Note that the performance metrics are only computed on positive image patches i.e., known to contain at least one helminth egg.

An overview of the results are shown in **Table 3**. The proposed approach showed a mean precision (P) of 98.4%, mean recall (R) of 80.9% and mean F-score (F) of 88.5% along all folders within the leave-one-out cross validation scheme.

**Table 3.**
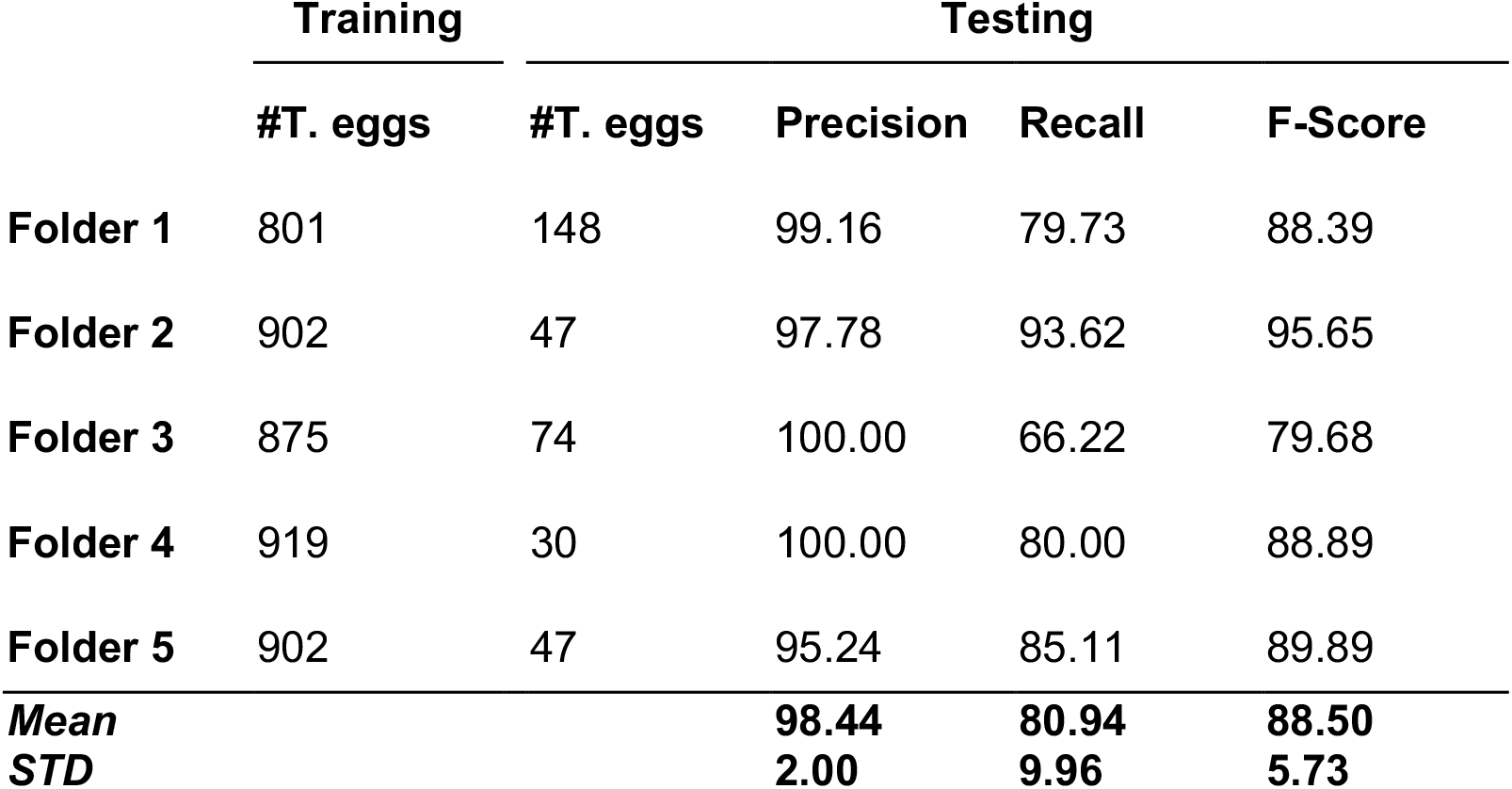
Detailed performance of the proposed methodology for *Trichuris* spp. detection using a leave-one-one cross validation scheme at the patient level. Note: ##T. eggs represent the number of *Trichuris* spp.

Additionally, we wanted to compare the results obtained with the proposed method, which is based on SSD and MobileNet networks, with a model based on FasterRCNN together with a ResNet50 backbone, a more complex and deeper network which also introduces the concept of residual connections. Our hypothesis is that, although this deeper network may have a higher districimantive power, it needs to be trained with a large number of training data. The network was trained with the same leave-one-out cross validation scheme as that used for training the proposed algorithm, and obtained a mean precision of 88.5% and a mean recall of 75.1%. It also should be noted that FasterRCNN-ResNet50 algorithm took much longer time to be trained and tested compared to the proposed SSD-MobileNet architecture, imposing a considerable computational cost. This results proves that the proposed method outperforms more complex networks, being an optimally cost effective architecture.

For further validation, we wanted to extend the analysis of the developed AI algorithm comparing their results at slide level to the ones obtained by all reference methods including both conventional microscopy-based analysis and manual remote analysis of digitized samples.

Digitized images from whole slides were divided into 512×512 pixels image patches using a sliding window procedure with an overlap of 64 pixels to ensure all helminth eggs are not cropped within images. All 27 slides coming from all subjects only positive for *Trichuris* spp. that were included in the analysis of Section 3.1 were analyzed by the AI algorithm, and Bland-Altman analysis between automated AI-assisted results and both reference methods were assessed.

Bland-Altman analysis showed a good agreement between AI-assisted results and both the reference method and the manual remote analysis, having a bias of −0.26, 95% confidence interval [−16.78,16.26] when compared to the results obtained with the conventional microscopy method, and a bias of −1.4 units, 95% confidence interval [−9.01,6.20] when compared to the remote analysis of digitized samples.

#### 3.2.2. Evaluation of the generalization capability

For further assessment of the proposed approach, we also wanted to test the generalization capability of the proposed methodology at detecting other helminths eggs than *Trichuris* spp. We extended our analysis by training the proposed method with all available subject samples, i.e. including those belonging to the one which were positive for a co-infection of *Trichuris* spp. and *Ascaris* spp. The training and evaluation of the extended version of the proposed method were based on a train-validation-test scheme. Of all available digitized slides from all subjects, 32 were randomly selected for training and validation, while the remaining 9 were used for testing. Both training (70%) and validation (15%) sets comprised 808 *Trichuris* spp. and 3649 *Ascaris* spp. eggs, while test test (15%) was composed by 136 *Trichuris* spp. and 647 *Ascaris* spp. eggs. Our interpretation is that the appearance of helminth eggs is independent across slides and therefore the training and testing may be done using images from the same subject as long as they belong to different slides.

Additionally, and in order to increase the discriminating power of the proposed approach and to better distinguish between both helminth eggs and all those structures that may be present on the images which are similar to objects under study, we deployed our trained model on all training slides, identified all false positives objects, labeled them as a separated class (namely artifact class) and constructed a multi-class model including three different classes, namely *Trichuris* spp., *Ascaris* spp. and artifacts. Thus, background objects which are difficult to discern and are confused with real helminth eggs were used as negative examples.

**Table 4** summarizes the results, and proves that the proposed model can be extended for the detection of different helminth eggs obtaining promising results including a mean precision of 94.36% and mean recall of 93.08%.

**Table 4.** Overview of the results obtained for the detection of both *Trichuris* spp. and *Ascaris* spp. helminths eggs.

|  | Precision | Recall | F-Score |
| --- | --- | --- | --- |
| ***Trichuris* spp.** | 95.31 | 89.71 | 92.43 |
| ***Ascaris* spp.** | 93.41 | 96.45 | 94.91 |
| **Mean** | 94.36 | 93.08 | 93.97 |

### 3.3. Built-in AI algorithms: operational deployment on technological platforms

The developed AI algorithm was integrated into both the telemedicine platform and the acquisition mobile app, enabling the operational deployment of the AI model so that it can be remotely used on demand by the telemedicine platform, or even could be executed during acquisition time through the mobile app. **Figure 4** shows the implementation of the AI algorithm into both technological platforms.

**Figure 4.**
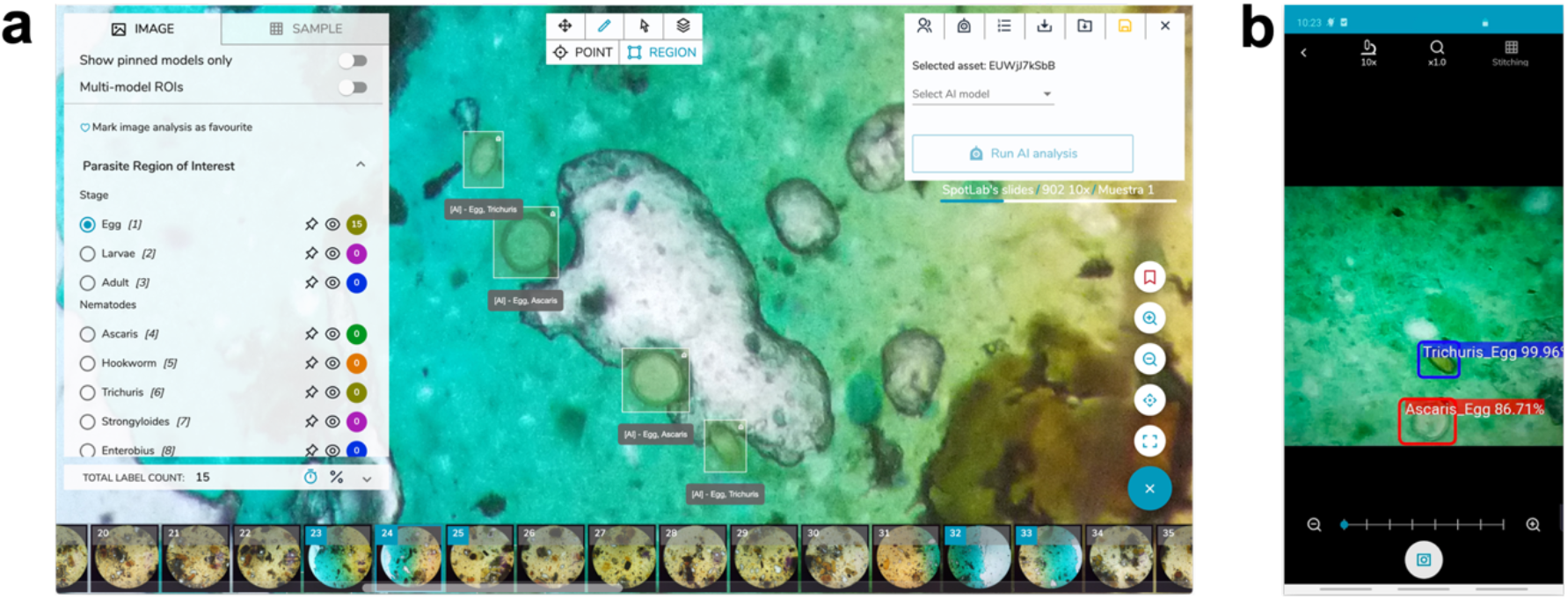
Operational deployment of the AI algorithm on technological platforms. **a** Deployment on the telemedicine platform. **b** Deployment on the mobile acquisition app.

The AI algorithm execution on the telemedicine platform is done on a cloud computing platform. Each time the images are uploaded, the system recognizes the type of sample based on the image metadata and executes the AI model which will detect the eggs and show them in a visualization environment which is accessed via the web. As an additional possibility of integration of AI models in the clinical workflow, we have embedded the AI model in the acquisition app. While the user examines the sample with the smartphone attached to the microscope objective, the smartphone processes the image acquired by the camera detecting the presence of eggs in real time. as the algorithm executes locally on the smartphone’s hardware. **Table 5** provides a report of the calculated time needed to perform the prediction for a single image patch of 512×512 pixels using different hardware configurations and technological platforms, as well as the average time needed to get a final prediction for a whole digitized image considering that each image is composed by 48 image patches.

**Table 5.** Comparison of the time needed to execute the AI algorithm with different technological platforms and hardware configurations.

| Tech. Platform | HW configuration | Patch image (s) | Whole Image (s) |
| --- | --- | --- | --- |
| Telemedicine platform | Low-moderate performance GPU (NVIDIA K80) | 0.04 | 1.97 |
| Telemedicine platform | Low-moderate performance CPU (Intel Xeon E5-2630 v3) | 0.20 | 9.60 |
| Mobile-phone | Low-moderate CPU (Snapdragon 820) | 0.25 | 12.00 |

## 4. Discussion and conclusions

In this work, we present an end-to-end pipeline for microscopy image digitalization and remote analysis together with novel deep learning algorithms for automatic assessment and quantification of parasitological infection of STH, mainly focused on the identification of *Trichuris* spp. eggs.

Microscopy remains the most widely used technique to complete a diagnosis of any of the NTDs, including STH which is one among the most prevalent diseases among this group. Visual diagnosis based on conventional microscopy is not very sensitive. It is also a time-consuming and subjective procedure, and requires specialised experts in the work field. Unfortunately, laboratory health workers density in STH-endemic areas is very low. Taking into account this limited resources together with the highly elevated number of patients in these areas, it is clear that the use of digital microscopes together with artificial intelligence algorithms for remote and automatic diagnosis of STH could constitute an advantageous tool. Although some systems have been previously proposed for the digitalization of images of STH, they require special hardware that has not been specifically designed for acquiring microscopy images (17) or might disrupt the usual laboratory workflow since they do not leverage conventional microscopes making it more complex to follow standard microscope diagnostics protocols (16). Our 3D-printed low-cost digitalization and image acquisition device was specially designed to not alter the daily routine in microscopy diagnosis laboratories. Additionally, the lack of primary care centers in high endemic areas of STH, entails a high need for field work to bring the diagnosis closer to remote areas. In this context, we designed a completely portable device enabling the digital diagnosis in those areas.

In this work we proposed the first complete pipeline from image digitalization using smartphone technology to remote analysis assisted by artificial intelligence methods.

The acquisition and digitization of the images are performed in a standardized manner and using a controlled procedure using a customized app which is directly linked to a telemedicine platform where the images were manually analyzed and tagged enabling the training of artificial intelligence algorithms.

The proposed pipeline was evaluated by comparing the quantification of *Trichuris* spp. infection with the reference method based on conventional microscope to both the remote analysis of digitized samples and the AI-assisted analysis y. The results showed a good agreement and minimal differences between methods when performing a global egg counting at slide level. These results proved and validated the use of the proposed digitalization and remote analysis pipeline as well as the proposed deep learning algorithms for the automatic identification of STH eggs.

Particularly, the proposed deep learning-based algorithm enables an automatic and objective identification of STH eggs. The method, which was trained and validated using a cross-validation scheme, achieved a relative high precision and recall results (98,44% and 80,94% respectively) for the identification and classification of *Trichuris* spp. eggs. We also wanted to compare the results with the ones obtained with more complex and deeper network architectures, and the results proved that the proposed relative simple architecture considerably reduces the computational cost while maintaining similar results.

Additionally, the results obtained suggest that remote and AI-assisted analysis of digitized images of Kato-Katz samples allows to detect in mean more eggs compared to the count using the conventional procedure, since the analysis can be performed in a more exhaustive manner. Moreover, these tools would potentially be useful for other use cases such as the evaluation of the effectiveness of MDA programmes.

For further validation and to illustrate the generalization capability of the method at identifying other helminth eggs than *Trichuri*s spp., we extended the analysis by training the algorithm including positive samples for both *Trichuris* spp. and *Ascaris* spp. The results (mean precision of 94,66% and mean recall of 93,08%) showed that the proposed method can be extended for the detection of different STH eggs although further validation work should be done in this direction. In particular the lack of presence of hookworm eggs in our samples, mainly caused by their tendency to disappear along time, may present a limitation of this study and should be included in future developments.

Finally, we proposed an operational implementation which allows to integrate the AI algorithm on both the remote analysis platform and the digitalization mobile app, opening a simple but potentially revolutionary use of the method on demand by invoking it through the telemedicine platform or in real time during image digitalization with the mobile app.

Next steps to scale the proposed system on the field require undertaking a large-scale clinical performance evaluation study to validate the entire pipeline and demonstrate its applicability where real time diagnosis is required.

Although there have been proposed effective and accurate molecular tools for the diagnosis of NTDs such as STH which are already implemented in developed countries (22), they may not be accessible in low- and middle-income countries. These molecular techniques such as quantitative PCR need expensive equipment and very well trained specialists. In order to achieve the goal established in the WHO 2030 roadmap (3), it is essential to have an effective and standardized diagnosis to accelerate the NTDs elimination. In this context, the proposed solution could reduce time, distances and expertise needed for microscopic analysis of helminth samples and therefore helping to make accurate STH diagnosis accessible.

## Authors Contribution

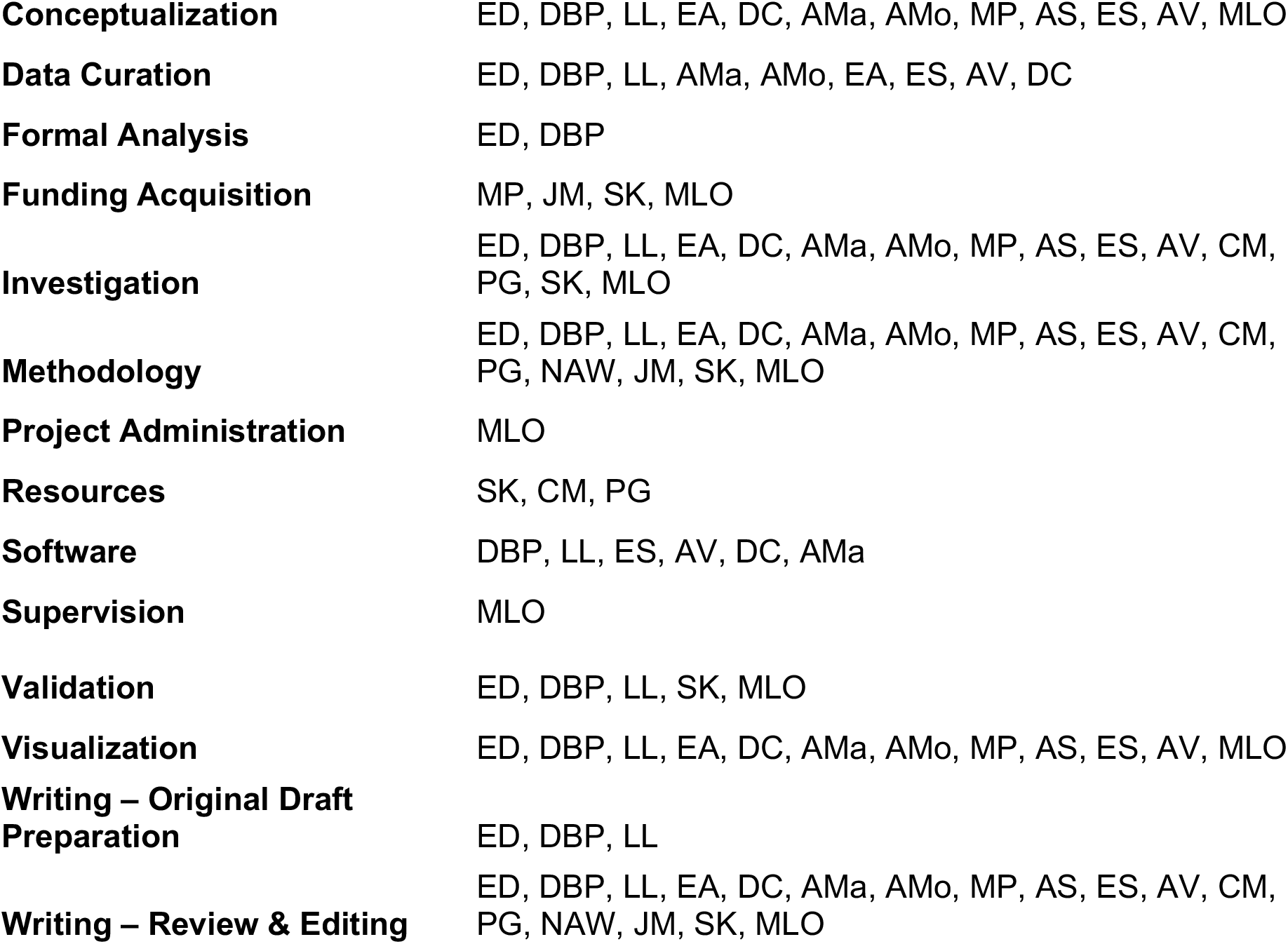

